# *Clostridioides difficile* stimulates *CCL20* expression in human colonoid monolayers in a transwell-based co-culture system that supports its anaerobic growth

**DOI:** 10.64898/2026.04.28.721417

**Authors:** Paola Zucchi, Adrianne D. Gladden, Andrew W. Day, Jules Dressler, Revathi Govind, Mohammad Almeqdadi, Jatin Roper, Albert Tai, Rebecca Batorsky, Carol A. Kumamoto

**Affiliations:** Department of Molecular Biology and Microbiology, Tufts University, Boston, Massachusetts, 02111, USA; Graduate School of Biomedical Sciences, Tufts University, Boston, Massachusetts, 02111, USA; Division of Biology, Kansas State University, Manhattan, KS 66506, USA; Department of Transplant Surgery, Tufts Medical Center, and Tufts University School of Medicine, Boston, MA 02111, USA; Department of Medicine, Duke University School of Medicine, Durham, NC 27710 USA; Department of Immunology and TUCF Genomics Core, Tufts University, Boston, Massachusetts, 02111, USA; Data Intensive Studies Center, Tufts University, Medford, MA, 02155 USA

**Keywords:** Colonoids/ *Clostridioides difficile*, toxins/ co-culture/ CCL20 Carol A. Kumamoto ORCID: 0000-0001-6352-4918

## Abstract

The pathogenic bacterium *Clostridioides difficile* is a major cause of antibiotic-associated diarrheal disease. Treatment of the disease is challenging because antibiotics used for treatment may also perpetuate the conditions that contributed to initial susceptibility. Elucidating the mechanisms of *C. difficile*/intestinal epithelium interaction is needed to facilitate the development of new therapeutic options. The studies described in this communication demonstrate the development of a tissue culture system that supported the growth of *C. difficile* in co-culture with a model of the human intestinal epithelium produced from colonoids, organoids derived from human colonic biopsies. Epithelial cell responses to *C. difficile* included upregulation of *CCL20*, encoding a chemokine that was previously shown to be upregulated in response to *C. difficile* challenge. Additionally, bacteria associated with the monolayer in a non-toxin dependent manner. This system will support future investigation of epithelium/*C. difficile* interactions during CDI and identification of mechanisms that drive pathogenesis by *C. difficile* in the human intestine.

## INTRODUCTION

*Clostridioides difficile* infection (CDI) is a leading cause of antibiotic-associated diarrheal disease and a major source of morbidity and mortality worldwide, particularly among hospitalized and elderly patients [1]. CDI most commonly occurs in individuals recently treated with antibiotics in part because antibiotics disrupt the gut microbiome, reducing the diversity and abundance of resident bacterial taxa that provide colonization resistance against *C. difficile* spore germination and vegetative growth.

Various approaches have been used therapeutically to treat this debilitating infection yet none are completely successful. The two primary antibiotics used for CDI treatment, vancomycin and fidaxomicin [2], are not 100% effective and many patients suffer from recurrence of infection after completion of treatment. Newer approaches include Fecal Microbiota Transplantation (FMT), in which an entire microbial community is transplanted into a patient, as an approach for preventing recurrent CDI [3]. However, rare adverse effects have been noted with FMT [4] prompting efforts to develop more targeted microbial interventions. In murine models, introduction of a single bacterium, *Clostridium scindens*, or a small consortium of bacteria reduced *C. difficile* disease [5, 6]. Feeding mice a diet high in plant polysaccharides reduced the ability of *C. difficile* to persist within the GI tract [7], suggesting that dietary approaches could contribute to lowering CDI incidence. Together, these observations highlight the continuing need for new and improved therapeutic approaches to combat CDI.

*C. difficile* is a strict anaerobe that persists in aerobic environments as a dormant spore. Following ingestion, these spores encounter germinants, such as the bile acid taurocholate, within the gastrointestinal (GI) tract [8]. Germination triggers the emergence of vegetative cells, which proliferate in the anaerobic parts of the GI tract and secrete glucosylating toxins, TcdA and TcdB [9], the primary virulence factors of *C. difficile*. Interactions between the organism, its toxins and the intestinal epithelium ultimately shape the progression and severity of disease.

One approach for characterizing *C. difficile*/epithelial interactions is to co-culture bacteria and epithelial cells in tissue culture. A variety of in vitro systems have been used previously to study the effects of *C. difficile* bacteria or its toxins [10–16]. Some studies have relied on specialized apparatus to generate anaerobic conditions to support *C. difficile* survival and growth; others have used bacterial components or short term culture with bacteria [17–20]. The goal of the present study was to develop a simple and accessible co-culture system that does not require specialized equipment yet supports robust growth of *C. difficile* alongside epithelial cells. To achieve this, we utilized organoids established from adult colonic biopsies, termed colonoids [21, 22]. Colonoids are derived from human stem cells and thus can be propagated in culture but are not transformed [23]. Colonoids can be differentiated in vitro and generate model epithelia that recapitulate many features of the human colonic epithelium.

Organoids in culture generate hypoxic environments [23]. We therefore tested the ability of an epidemic strain of *C. difficile* to grow in co-culture with a differentiated colonoid monolayer produced on a transwell and incubated in an aerobic atmosphere. Transwells were used so that the apical surface of the differentiated monolayer was accessible for bacterial interaction. Results showed that bacteria proliferated in this system and maintained viability for at least 20 hrs. We detected association of bacteria with the monolayer that increased over time. The epithelial cells responded to bacterial toxin production by upregulating the chemokine CCL20, reproducing a response that has been previously demonstrated [24]. This system provides a simple approach that will enable further investigation of *C. difficile*/epithelial cell interaction.

## MATERIALS AND METHODS

### Bacterial growth and spore preparation

*C. difficile* strain UK1 (a NAP1/027/BI human epidemic strain [25]) was used for all experiments except as noted. For studies of nontoxigenic *C. difficile,* R20291 (a NAP1/027/BI human epidemic strain closely related to UK1 [26]) and an R20291::*tcdR* mutant strain [27] were used. Lack of TcdA/TcdB production by the *tcdR* mutant strain was verified by testing culture supernatants with the tgcBIOMICS Clostridium difficile Toxin A or B ELISA kit (TGC-E002-1).

Bacteria were cultured on prereduced BHIS medium at 37°C in an anaerobic chamber. For sporulation, cultures were grown in prereduced TY broth and plated on prereduced 7030 medium [28]. After one week, bacterial growth was collected from the plates. For some experiments, the suspension was incubated in water at 4°C overnight. Spores were collected by centrifugation at 15000 rpm for 15 min in a Hitachi R20A2 rotor. Spores were washed with water and purified by centrifugation in a discontinuous Histodenz (Sigma D2158) gradient as described previously [29]. Purified spores were washed with water, aliquoted and stored at - 20°C.

For infections, spores were pre-germinated by treatment with germinants under aerobic conditions followed by addition to the colonoid monolayer system where vegetative cells would outgrow when anaerobic conditions were encountered. A spore aliquot was thawed, diluted to approximately 5 x 10^7^ spores/ml and incubated at 60°C for 10 min. Spores were diluted with an equal volume of 120 mM glycine (Sigma G7126), 4 mM sodium cholate (Sigma C9282) in PBS and incubated at 37°C for 15-30 min to produce pre-germinated spores. The pre-germinated spores were either added directly to epithelial cell monolayer cultures or were collected by centrifugation (Eppendorf 5415 centrifuge, 13200 rpm, 5 min) and resuspended in the same volume of PBS before addition to monolayer cultures. 10 ul of pre-germinated spores (approximately 5 x 10^5^ spores) were added per 6.5mm diameter transwell (Costar 3413).

### Preparation of colonoid monolayers on transwells

Colonoid line CJ50 was derived from colonic biopsies obtained from a 67 year old Chinese female undergoing routine outpatient colonoscopy for colorectal cancer screening at Tufts Medical Center (IRB #11652, principal investigator: Jatin Roper, date of approval 8/18/2025). The cells were propagated in Matrigel (Corning 356231) as described previously [30]. Medium composition was as described [22] except that L-WRN conditioned medium (L-WRN cells ATCC CRL- 3276 [31]) was used at 65% instead of L-WNT3A, R-Spondin and Noggin conditioned media, Primocin (InvivoGen ant-pm-1) was used instead of pen-strep and 5 uM Y27632 (Rho kinase inhibitor, Sigma Y0503) was added for the first 48 hrs of culturing.

For preparation of monolayers on transwells, colonoid spheres were recovered from Matrigel, washed with PBS and digested with TrypLE Express (Gibco 12605-010) for 2.5 min at 37°C. After digestion, cells were diluted with 1 volume advanced DMEM (Invitrogen 12634-028), 1% Glutamax (Gibco 35050-061), 20% FBS (Fisher 26140079) and filtered through a 40 um cell strainer. Cells were diluted with 0.4% Trypan blue, counted with a Corning CytoSmart counter and diluted to 2.5 x 10^6^ cells/ml with warmed 65% LWRN medium containing Y27632.

Matrigel was diluted with CMGF- (advanced DMEM medium containing additives but not growth factors [22]) to 33 or 66 ug protein/mL, 125 ul was applied to each 6.5mm diameter, 0.4 um pore, polycarbonate filter transwell (Costar 3413) and incubated at 37°C for 2 hours. Liquid was removed from the transwell, the basolateral compartment was filled with 0.5 ml 65% medium and 50 ul of medium was added to the apical compartment. Cells (2.5 x 10^5^ cells per 6.5 mm diameter transwell) were added and the seeded transwells were incubated in a tissue culture incubator with 5% CO2 at 37°C.

The next day, transwells were given fresh 65% L-WRN medium containing 5 uM Y27632 (Sigma Y0503), replacing the basolateral medium and adding 0.2 ml to the apical compartment. The next day, the medium was changed to 65% without Y27632 (0.4 ml apically and 0.5 ml basolaterally) and was changed again after 2 days when the monolayer was typically confluent.

Confluent monolayers were differentiated by washing with CMGF-, followed by feeding with Differentiation Medium (DM) [22, 32] containing Primocin. Medium was changed the next day with DM containing Primocin. Monolayers were fed 2 days later (day 3) (DM without Primocin) and in the late afternoon of day 4 or in the morning of day 5 (DM without Primocin). Monolayers were inoculated with pre-germinated spores on day 5.

For measurement of trans-epithelial electrical resistance (TEER), a Millicell ERS-2 volt/ohm meter was used following manufacturer’s protocol.

### Co-culturing *C. difficile* with colonoid monolayers

Pre-germinated spores (approximately 5 x 10^7^ per ml) were prepared as above and 10ul of the preparation was added gently to the apical medium of the transwell. Control samples were inoculated with 10ul of buffer. The plates were wrapped with Parafilm and incubated in a standard tissue culture incubator (37°C, 5% CO2, room air).

At various times, transwells were taken into the anaerobic chamber (2 transwells at a time) to estimate the number of bacteria per transwell. The first 170 ul of apical medium was carefully removed. This medium was not turbid and was discarded. The second approximately 170 ul of medium was collected into the pipet tip and used to wash the monolayer twice, then collected into a pre-weighed tube. For inoculated transwells, this sample was typically turbid. 10 ul of this sample was immediately diluted with 1 ml of pre-reduced PBS to measure CFU. The remainder of the sample was removed from the chamber and weighed to determine the volume recovered from each transwell after correction for the 10 ul removed for CFU.

Transwell monolayers were removed from the chamber and fixed with 4% paraformaldehyde for 15 min at room temp, followed by washing 3 times with PBS at room temperature. Transwells were stored at 4°C in PBS.

For some experiments, transwells were inoculated with vegetative *C. difficile* cells. Strains were grown in pre-reduced TY broth and harvested during log phase. A sample of each culture was diluted 1/50 into pre-reduced PBS. DM medium was pre-reduced and 0.4 ml was transferred to a gasketed screw cap tube for each inoculation. 10 ul of bacteria diluted in PBS was added to each tube (or PBS only for mock inoculation). The caps were screwed on tightly and tubes removed from the chamber. The transwells were inoculated by removing their apical medium and replacing it with the bacteria-containing medium. Transwells were inoculated with approximately 5 x 10^4^ CFU.

### Detection of secreted protein production by colonoid cells

In some cases, after incubation of monolayers with *C. difficile* for 16 hours, the cultures were treated with brefeldin A (Sigma B6542) (25 ug/ml final added to the apical and basolateral medium) for 4 hours. This treatment was used to block protein secretion and facilitate detection of the secreted protein CCL20.

### Preparation of anti *C. difficile* antiserum

UK1 bacterial cells were grown to exponential phase in TY broth. Cells were washed three times with PBS, resuspended in 2% (vol/vol) neutral buffered formalin and incubated overnight at room temperature. Cells were washed extensively with PBS and resuspended at a concentration of 4 x 10^9^/ml. Anti-UK1 antisera were raised in rats by Pocono Rabbit Farm and Laboratory.

### Staining and imaging of monolayers

Membranes were cut from the transwell with a scalpel and permeabilized in 0.1% Triton X-100 in PBS for 15 min. Membranes were washed 3 times with PBS and blocked by incubation in 2% BSA, 1% normal donkey serum, PBS for 1-2 hours. Probing with primary antibodies in 2% BSA using manufacturer recommended dilutions was conducted at 4°C overnight. Primary antibodies: anti-NHE3 (Novus NBP1-82574, 1:100), anti-MUC2 (GeneTex GTX100664, 1:200), anti-β-catenin (Invitrogen 13-8400, 1:100), anti-CCL20 (Invitrogen PA5-47517, 1:20), anti-formalin fixed UK1. Membranes were washed 3 times with PBS, and probed with secondary antibodies (Donkey anti-rat Alexa Fluor 488 (Invitrogen A21208, 1/1000), Donkey anti-rabbit Alexa Fluor 488 (Invitrogen A21206, 1/1000), Donkey anti-rabbit Alexa Fluor 594 (Invitrogen A21207, 1/1000), Donkey anti-mouse Alexa Fluor 594 (Invitrogen A21203, 1/1000), or Donkey anti-goat Alexa Fluor 594 (Invitrogen A32758, 1/1000)) in 2% BSA for 1-2 hours. Membranes were washed with PBS containing Hoechst 33342 (Invitrogen H3570, 1/1000) and for some experiments, phalloidin-Alexa Fluor 680 (Invitrogen A22286, 1/1000).

Membranes were imaged using a Leica DMi8 with a 63X oil immersion objective. Images were collected using LasX software. Image analysis and figure construction was performed using Fiji. Images shown of the XY plane are maximum intensity Z projections. Orthogonal views were analyzed using LasX. Bacteria were defined as in a vertical orientation if the long axis of the bacterium was perpendicular to the long axis of the monolayer (90° ± 45°) in both the XZ and YZ planes.

### Single cell RNA Sequencing (scRNASeq)

Monolayers were cultured on 12 mm diameter 0.4 um polycarbonate filters (Millicell PIHP01250) and differentiated as described above. Spores were treated with glycine and sodium cholate, as above, collected by centrifugation (Eppendorf 5415, 13200 rpm, 5 min) resuspended in PBS and 1 x 10^5^ spores in 10 ul or PBS was added to the apical medium. Transwells were incubated as described above.

After 4, 15, 20 or 25 hours of incubation, monolayers were washed with 100 ul PBS, EDTA 0.5 mM, pH 8 and incubated with TrypLE Express, 600 ul in the basolateral and 400 ul in the apical compartments at 37°C for 12 min.

One hundred ul 50% FBS was added to the apical compartment and the cells were suspended by pipeting up and down several times. Cells were passed through a 40 um cell strainer and recovered in tubes precoated with 0.1% BSA. The transwells were then washed with 400 ul CMGF-, which was passed through the strainer and combined with the previous filtrate. Cells were collected by centrifugation at 300g x 5 min. Most of the supernatant was gently removed. Cells were resuspended in 1 ml of 0.04% BSA, 5 uM Y27632 and collected as above. Using a wide bore tip, the pellet was resuspended in the remaining supernatant by pipetting up and down. Cells were counted with a Corning Cytosmart counter and adjusted to 1 x 10^6^ cells/ml.

The remaining steps were performed following manufacturer’s protocol. Briefly, the cell suspension was loaded along with reagents, barcoded Single Cell 3’ v3.1 Gel Beads and Partitioning Oil onto the Chromium Next GEM Chip G. Cells were partitioned into GEMS using the Chromium Controller. Gel beads were then dissolved, releasing primers, and the cells were lysed. Reverse transcription was performed to generate cDNA tagged with cell specific barcodes and Illumina TruSeq Read 1 sequencing primer. GEMS were broken and first strand cDNA was purified and PCR amplified. After enzymatic fragmentation followed by size selection, the cDNA was PCR amplified, incorporating a sample index and primers P5, P7, and TruSeq Read 2 for Illumina sequencing. Library quality was assessed using an Agilent Bioanalyzer. Paired end sequencing was performed by the Tufts University Genomics Core Facility.

### scRNA-Seq data analysis

Initial data analysis including demultiplexing, barcode processing and aligning reads to the human genome was performed using Cell Ranger. Seurat version 5.0.2 [33] was used to analyze the scRNA-seq dataset. Transcriptomes with less than 250 or more than 5000 detected expressed genes or more than 40% mitochondrial expressed genes were removed from the analysis. The *C. difficile* toxins TcdA and TcdB are cytotoxic and we therefore expected that some responding cells would be unhealthy. We analyzed cells with up to 40% mitochondrial expressed genes to retain some of the less healthy cells. Gene expression profiles of two replicates of Mock and CDI colonoid monolayers were log normalized, scaled and subjected to dimensionality reduction with PCA (30 PCs). Results were integrated using Harmony integration (30 PCs) to mitigate sample variability and group transcriptionally similar cells together.

Clustering using the Louvain algorithm was conducted at resolution 0.1, yielding 4 clusters. Cluster markers were identified using FindAllMarkers() with logfc.threshold = 0.25, min.pct = 0.1, only.pos = TRUE to compare gene expression in cells of one cluster to all other cells in the other 3 clusters. The markers obtained were compared to previously described markers for colonic cell types [34] from the Gene Set Enrichment Analysis (GSEA) Human Molecular Signatures Database. SeuratExtend was used to produce dot plots [35].

Cell type annotation was conducted using SingleR [36], a reference-based method for annotating cells that compares individual cell transcriptional profiles with the transcriptomes of reference cells. Colonoid cells were assigned the identity of the reference cell that was most comparable using the human colon reference published by Smillie et al.[37], that included 12 healthy colon biopsy samples. After annotation, cell type counts were exported for analysis in Excel. Related cell types were combined into groups to simplify the findings. The following labels were used: Enterocytes = Enterocytes + Immature Enterocytes 1 + Immature Enterocytes 2; Goblets = Goblets + Immature Goblets; M-like cells = M cells; Other = Stem + TA 1 + TA2 + Tuft + NA. Several cell types were grouped into the Other category because of their very low abundance. Graphpad Prism was used to produce graphs of cell type composition.

Differentially expressed genes upregulated or downregulated in *C. difficile* co-cultures vs mock samples were identified for each cluster using the Seurat FindMarkers() function (logfc.threshold= 0.25, min.pct = 0.10).

### RNA extraction and real time RT qPCR

Colonoid cells (4-5 x 10^5^ cells per transwell) were seeded on 12 mm diameter 0.4 um polycarbonate filters and incubated for 5 days. Medium was changed to differentiation medium and the transwells were incubated for another 5 days. Primocin was withdrawn 2 days before inoculation with pre-germinated UK1 spores.

Pre-germinated UK1 spores (1 x 10^6^) or buffer were added to each transwell. After incubation for 16 hrs, the transwell was moved to an empty well, apical medium was removed and the membrane was washed. Trizol (0.3 ml) was added to the apical compartment, pipeted up and down 5 times and transferred to a tube. A second 0.3 ml of trizol was added and allowed to incubate briefly. After pipetting up and down, the second trizol extract was combined with the first and frozen at-80°C. RNA was extracted using the Purelink RNA mini kit (Invitrogen 12183025) with on column DNase treatment (Invitrogen 12185-010) following manufacturer’s protocol. An aliquot of RNA was used to generate cDNA by reverse transcription with Superscript III (Invitrogen 18080044) following manufacturer’s protocol. cDNA was diluted 1:3 and used as template for RT PCR reactions.

*CCL20, ANKRD37* and *GAPDH* gene expression was measured in cDNA by qPCR using 2x SYBR Green MasterMix (AppliedBiosystems 4334973). qPCR reactions were run on a Roche LightCycler 480 II as described [38]. *CCL20* and *ANKRD37* primer sequences (from Harvard PrimerBank) were: hCCL20 F1 5’ TGCTGTACCAAGAGTTTGCTC 3’; hCCL20 R1 5’ CGCACACAGACAACTTTTTCTTT 3’; hANKRD37 F1, 5’ TTAGGAGAAGCTCCACTACACAA 3’; hANKRD37 R1, 5’CACTGGCTACAAGCAGGCT 3’. *GAPDH* primer sequences were designed using Primer BLAST: hGAPDH F1, 5’ CATGTTCGTCATGGGTGTGAA 3’ and hGAPDH R1, 5’ GACTGTGGTCATGAGTCCTTCC 3’. Primer specificity was demonstrated by Sanger sequencing of the qPCR product. No template controls yielded no signal. Gene expression relative to the average of mock inoculated samples was calculated by the 2^-ΔΔCT^ method. [39].

## RESULTS

### A model of the human intestinal epithelium derived from human colonoids

Differentiated human colonoid monolayers were used as a model of the intestinal epithelium. To produce monolayers for co-culture with bacteria, the colonoids were enzymatically disaggregated and seeded onto Matrigel-coated polycarbonate Transwell filters. The monolayers were grown to confluence and then incubated in differentiation medium. Immunofluorescent staining of fixed, differentiated monolayers showed that markers of differentiated epithelial cells such as NHE3 (sodium-proton exchanger, marker of colonocytes; Fig. 1 B,C) and MUC2 (mucin, marker of goblet cells; Fig. 1, E,F) were expressed. The cells were polarized with nuclei localized towards the basal side (Fig. 1 J-L). Fig. 1 C, F overlays show the apical side of the monolayer with the basal side below. Apical markers therefore obscure the nuclei in some regions of the images. Further, the cells formed intercellular junctions (detected by β-catenin staining of adherens junctions, Fig. 1 H,I) and trans-epithelial electrical resistance values ranging from 300-450 Ω cm^2^ were typically obtained (Fig 1 M). This model of the human intestinal epithelium thus recapitulated many features of the human colonic epithelium.

**Figure 1.**
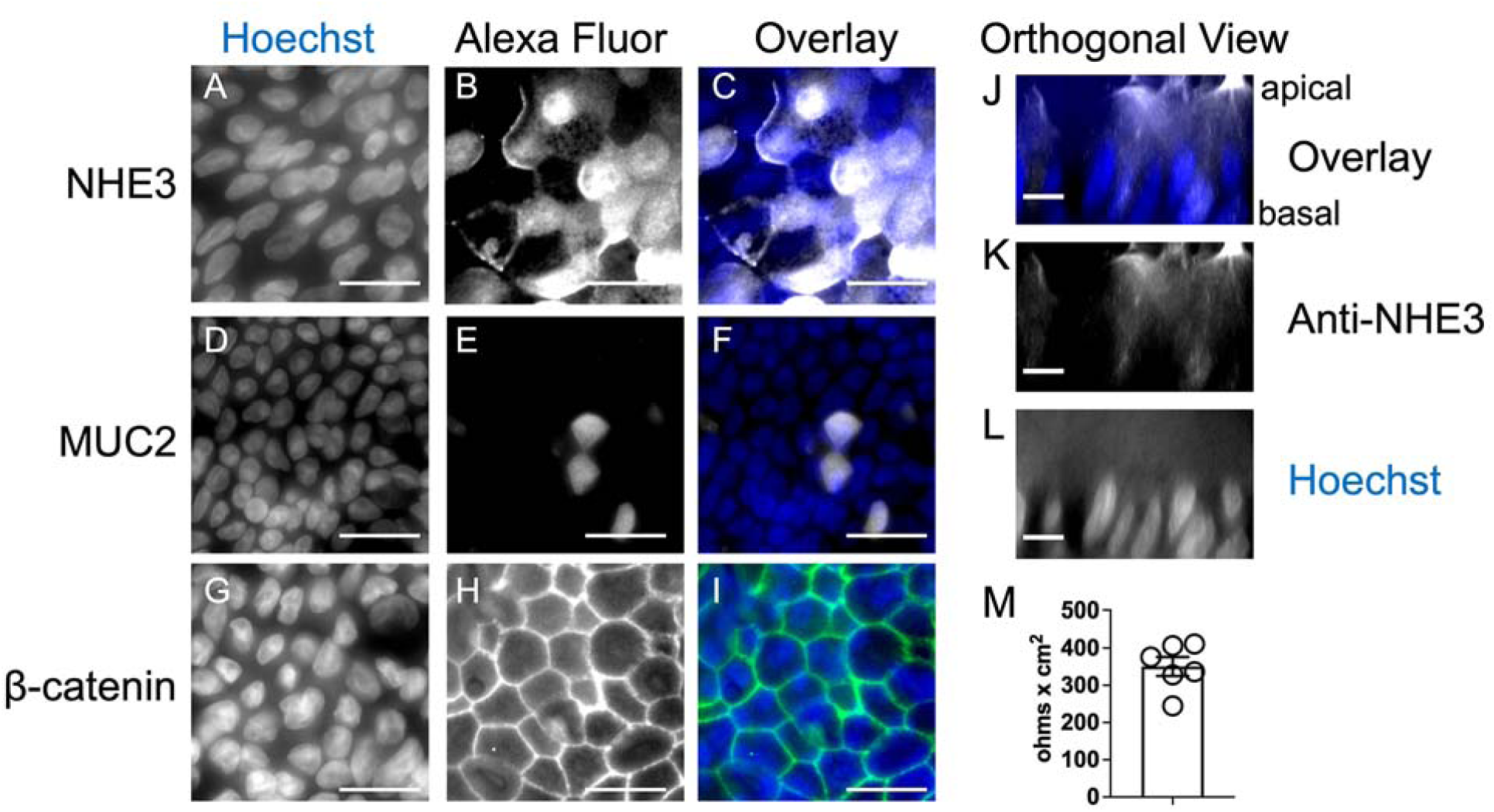
Colonoids differentiated on polycarbonate filters in transwells produce monolayers of polarized, differentiated cells with barrier function. Colonoids were cultured in Matrigel, harvested, dissociated and seeded on polycarbonate filters in Transwells. After growth to confluency and incubation in differentiation medium for 5 days, the monolayers were fixed with 4% PFA. Fixed monolayers were permeabilized and stained with antibody and Hoechst as described in Materials and Methods. Immunofluorescent signal was detected using a Leica DMi8 microscope and visualized with a 63x oil immersion objective. Panels A-I: staining with Hoechst (A, C (blue), D, F (blue), G, I (blue)); staining with anti-NHE3 (B, C (white)); staining with anti-MUC2 (E, F (white)); staining with anti-β-catenin (H, I (green)). Scale bar, 20 um. Panels J-L: Orthogonal views of the Z plane showing polarization. Staining with Hoechst (J,L (blue)); staining with anti-NHE3 (K,L (white)). Scale bar, 10 um. Panel M: Transepithelial electrical resistance was measured as described in Materials and Methods. Each symbol shows results from a different monolayer, bar shows the mean with standard error of the mean.

### Human colonoid monolayers support the growth of the anaerobic pathogen *C. difficile*

We tested whether this model colonic epithelium would support growth of the anaerobic bacterium *C. difficile* in co-culture in aerobic conditions. Monolayers were incubated without antibiotics before inoculation with *C. difficile*. Spores of *C. difficile* strain UK1 [25] were pre-germinated and added to the apical compartment of the transwell (5 x 10^5^ spores/transwell). Mock inoculated cultures received buffer. The co-cultures were incubated in a standard tissue culture incubator with a 5% CO_2_, room air atmosphere.

After incubation, the number of viable *C. difficile* bacteria in the co-cultures was estimated by sampling the apical medium of the transwell. The inoculated transwells were transferred to the anaerobic chamber and the most apical medium, expected to be more aerobic, was removed and discarded; this sample was not turbid, consistent with low bacterial density. The lower apical medium, next to the monolayer, was carefully collected into a pipet tip and used to wash the monolayer gently two times; this sample was typically turbid, consistent with higher bacterial density. The apical culture medium was transferred into a pre-reduced, pre-weighed Eppendorf tube and a sample was immediately diluted with pre-reduced PBS and further diluted for plating on pre-reduced BHIS plates. Colonies were enumerated and total CFU/transwell was calculated. Results showed an increase in CFU per transwell after 4 hours of co-culture (4 hrs, Fig. 2A) or 20 hours of co-culture (20 hrs; Fig. 2A) compared to the inoculum (inoc; Fig. 2A). *C. difficile* bacteria thus grew and survived in co-culture with colonoid monolayers despite the aerobic atmosphere of the incubator.

**Figure 2.**
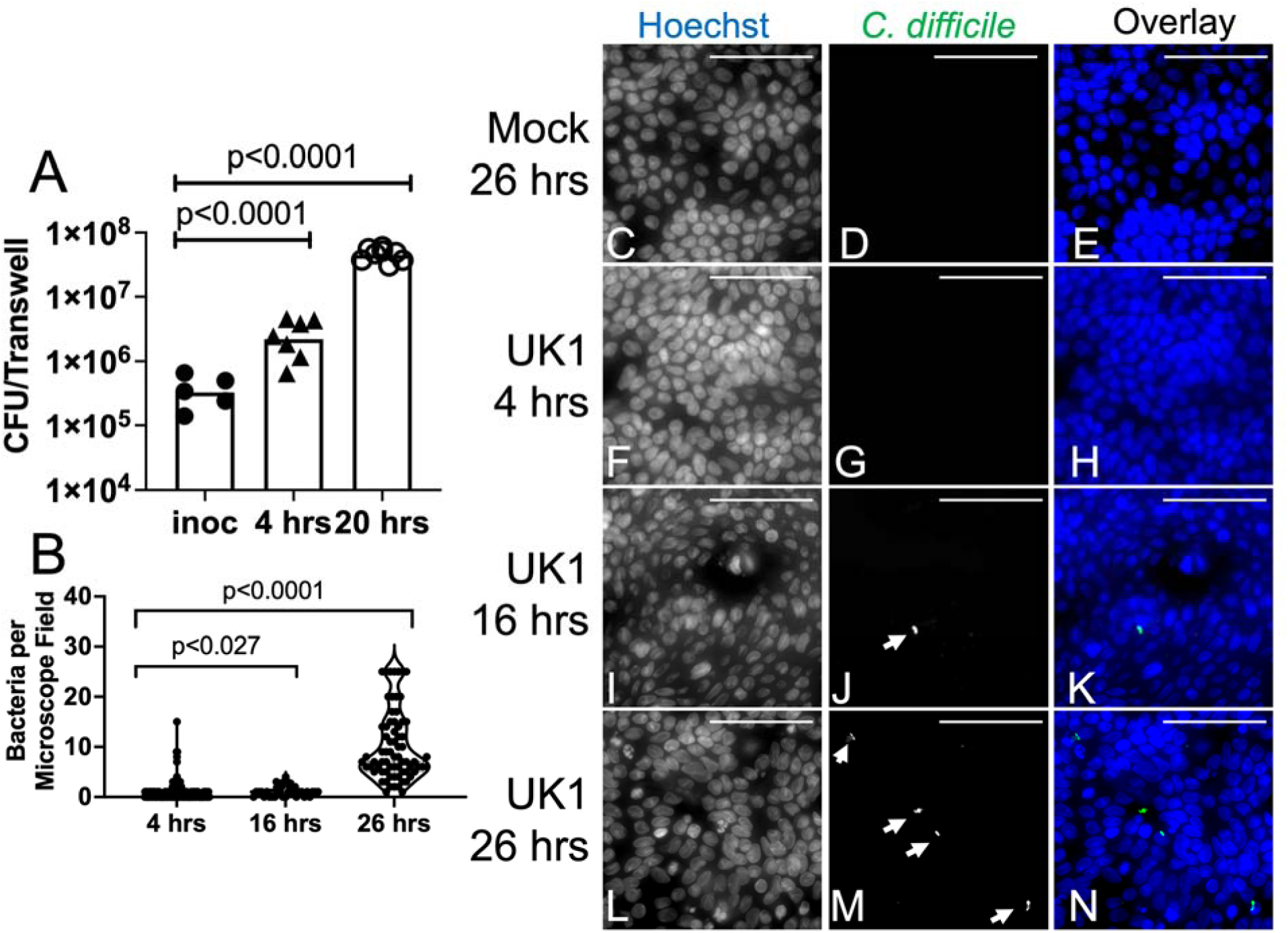
Differentiated colonoid monolayers support growth of C. difficile. Differentiated colonoid monolayers were produced as in Figure 1. Panel A: Monolayers were inoculated with pre-germinated spores of *C. difficile* strain UK1. Total CFU/transwell detected in the apical culture medium is shown. Each symbol shows results from a different transwell. Bar, geometric mean. p values, one-way ANOVA performed with log transformed data, using Dunnett’s multiple comparison test. Composite results from 3 experiments. Panel B: Bacteria per microscope field were scored visually and counted. Each symbol shows results from one 63x microscope field. 4 hrs, n=201 fields, median=0; 16 hrs, n=34 fields, median=1; 26 hrs, n=71 fields, median =7; combined results from 2-3 experiments. p, Kruskal Wallis with Dunn’s multiple comparisons test. Panels C-N: Monolayers were inoculated with pre-germinated spores of *C. difficile* strain UK1 or buffer (Mock). Monolayers were incubated for 4 hours (panels F-H), 16 hours (panels I-K) or 26 hours (panels C-E, L-N) after inoculation. After removing the apical medium and washing the apical surface as described in Materials and Methods, epithelial monolayers were fixed in 4% PFA and stained for immunofluorescence. Stained with Hoechst (C, E (blue), F, H (blue), I, K (blue), L, N (blue)); stained with anti-*C. difficile* (D, E (green), G, H (green), J, K (green), M, N (green)). White arrows indicate bacteria. Scale bar, 50 um.

*C. difficile* associates with the epithelium and exhibits vertical orientation.

In addition to proliferating, bacteria became associated with the monolayer during co-culture. After monolayers were washed as described above for enumeration of *C. difficile* CFU, the transwells were removed from the anaerobic chamber and fixed with 4% paraformaldehyde, followed by washing with PBS. The polycarbonate filter was cut from the transwell and incubated for immunofluorescence as described in Materials and Methods. Filters were stained with anti-*C. difficile* antiserum, Hoechst and anti-NHE3. The number of bacteria associated with the colonoid monolayers was scored. After 4 hrs of co-culture, the median number of bacteria per 63x field was 0, after 16 hrs, the median was 1 and after 26 hrs, the median was 7 (Fig. 2B-N; bacteria marked with white arrows). No bacteria were detected in association with mock inoculated monolayers. (Fig. 2C-E). These results show that bacteria associated with the monolayer and their number increased over time.

Orthogonal views of the colonoid images demonstrated that the majority of monolayer-associated *C. difficile* exhibited a vertical orientation. Images of the washed monolayers used in Figure 2 were analyzed in the XZ and YZ planes and bacterial orientation was scored visually. A bacterium was defined as vertical if its long axis was at a 90° (± 45°) angle to the long axis of the monolayer in both XZ and YZ planes. Fig. 3 A-D shows a vertically-oriented bacterium in relation to the NHE3-stained apical membrane of the monolayer. Quantification of vertical and non-vertical bacteria after 4, 16 or 26 hours of co-culture showed significant enrichment for vertical bacteria at all time points (Fig. 3E, Fisher’s exact test versus 50:50 distribution of bacteria).

**Figure 3.**
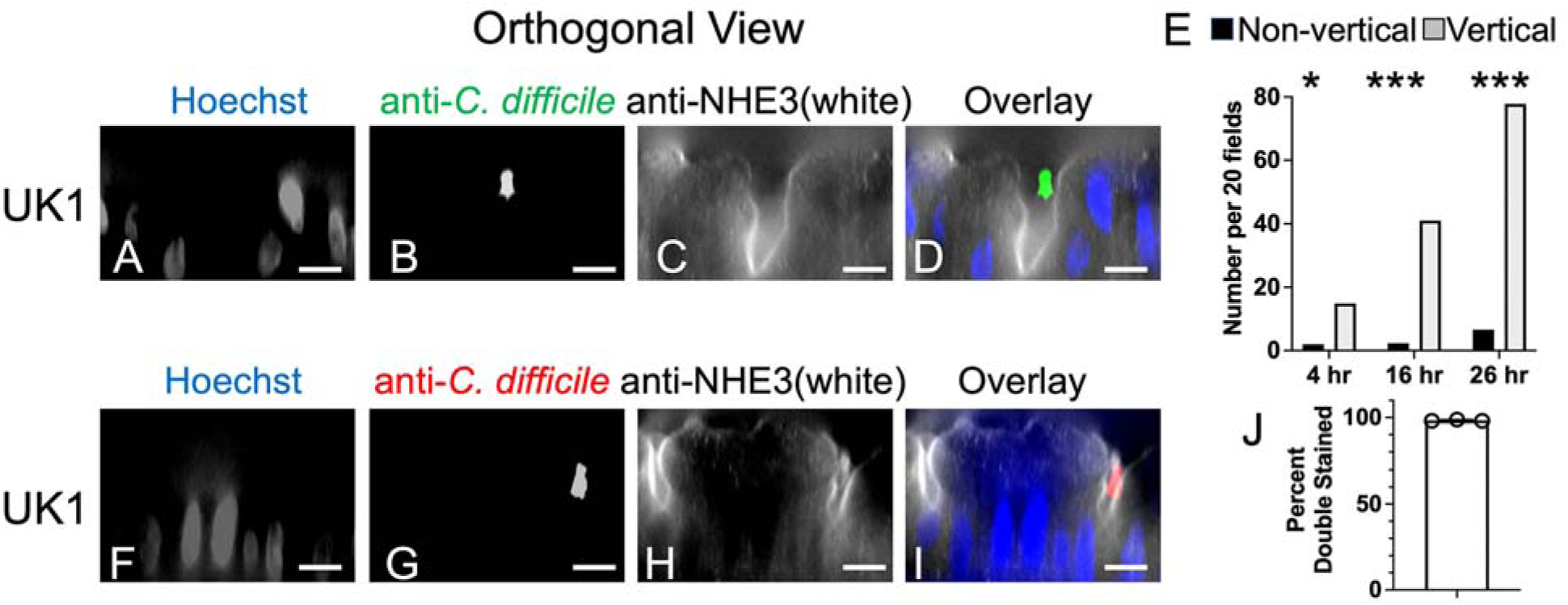
Monolayer-associated *C. difficile* bacteria were extracellular and in a vertical orientation relative to the monolayer. Differentiated colonoid monolayers were produced as in Figure 1. Panels A-D: Monolayers were co-cultured with pre-germinated UK1 spores for 16 hrs, washed, fixed with 4% PFA, permeabilized and stained. Orthogonal views are shown. Apical surface is above, basal surface below. Stained with Hoechst (A, D (blue)); stained with anti-*C. difficile* (B, D (green)); stained with anti-NHE3 (C, D (white)). Scale bar = 10 um. Panel E: Bacterial orientation was scored visually. A vertical bacterium was defined as a bacterium with the long axis at a 90° (± 45°) angle relative to the long axis of the monolayer in both the XZ and YZ planes. 17-19 63x microscope fields were scored and numbers were normalized to 20 fields. Grey bars show vertical bacteria; black bars show non-vertical bacteria. Results tested against a 50:50 distribution by Fisher’s exact test. *, p < 0.03; ***, p<0.0001. Panels F-I: Monolayers inoculated with pre-germinated spores of strain UK1 were incubated for 20 hours, washed and fixed with 4% PFA. Before permeabilization, bacteria were stained with anti-*C. difficile* primary antibody and Alexa Fluor 594 conjugated secondary antibody (red). Monolayers were then permeabilized with 0.1% Triton X-100 and stained with anti-*C. difficile* primary antibody and Alexa Fluor 488 conjugated secondary antibody (green). Stained with Hoechst (F, I (blue); stained with anti-*C. difficile* prior to permeabilization (G, I (red)); stained with anti-NHE3 (H, I (white)). Scale bar = 10 um. Panel J: Bacterial staining was scored visually (60-150 bacteria scored per experiment; 3 experiments) and the percent of bacteria that were double stained was plotted. Over 98% of bacteria were double labeled. Rare bacteria that stained only after permeabilization (green only) were detected. No bacteria were labeled with red only.

Bacterial accessibility to antibody in the absence of epithelial permeabilization was analyzed to test whether monolayer-associated bacteria were extracellular. Monolayers were co-cultured with pre-germinated UK1 spores for 20 hrs. Washed, fixed monolayers were stained with anti-*C. difficile* without permeabilization, followed by incubation with Alexa Fluor 594 conjugated secondary antibody (red; Fig. 3 G, I). The monolayers were then permeabilized and stained with anti-*C. difficile*, followed by incubation with Alexa Fluor 488 conjugated antibody (green). Most of the bacteria were double-labeled (98%, Fig. 3J), indicating accessibility to antibody without permeabilization. Rare bacteria were stained only green. In these cases, a nearby bacterium appeared to be interfering with staining. No bacteria stained only red. These results show that monolayer-associated bacteria were typically found vertically oriented in an extracellular location.

### Identification of CCL20 as a marker of early colonoid responses to *C. difficile* challenge

We conducted a single cell RNASeq (scRNASeq) study of colonoids in co-culture with *C. difficile* or with buffer only (referred to as Mock). A more detailed analysis of the responses of colonoids to *C. difficile* challenge over a time course will be presented elsewhere. Our analysis here focused on identifying gene markers that could be used to monitor the response of colonoids to *C. difficile* in co-culture.

Differentiated colonoid monolayers were produced as described for Fig. 1 and inoculated with pre-germinated spores of strain UK1 (or buffer). Two replicates of the experiment comparing a Cd co-culture with a mock culture after 15 hrs of incubation were performed, yielding 4 samples: Mock replicate a (Ma), Mock replicate b (Mb) *C. difficile* co-culture replicate a (CDa) and *C. difficile* co-culture replicate b (CDb).

As described in Materials and Methods, single cell suspensions were produced and processed for scRNASeq using the 10x platform. Sequencing was conducted by the Tufts University Genomics Core Facility. Sequences were demultiplexed and aligned to the human genome using Cell Ranger. Results were analyzed using Seurat 5.0.2 [33] in R. Transcriptomes with less than 250 or more than 5000 detected expressed genes or more than 40% mitochondrial expressed genes were removed from the analysis. Transcriptomes with up to 40% mitochondrial expressed genes were included because *C. difficile* toxins are cytotoxic and some responding cells were expected to show signs of damage.

Transcriptomes were processed as described in Materials and Methods, integrated using Harmony integration and displayed in a UMAP based on their transcriptional similarity. Clustering using the Louvain algorithm yielded 4 clusters (Fig. 4A). Transcriptomes with high mitochondrial content (up to 40%) were found in multiple clusters (Supplemental Figure S1).

**Figure 4.**
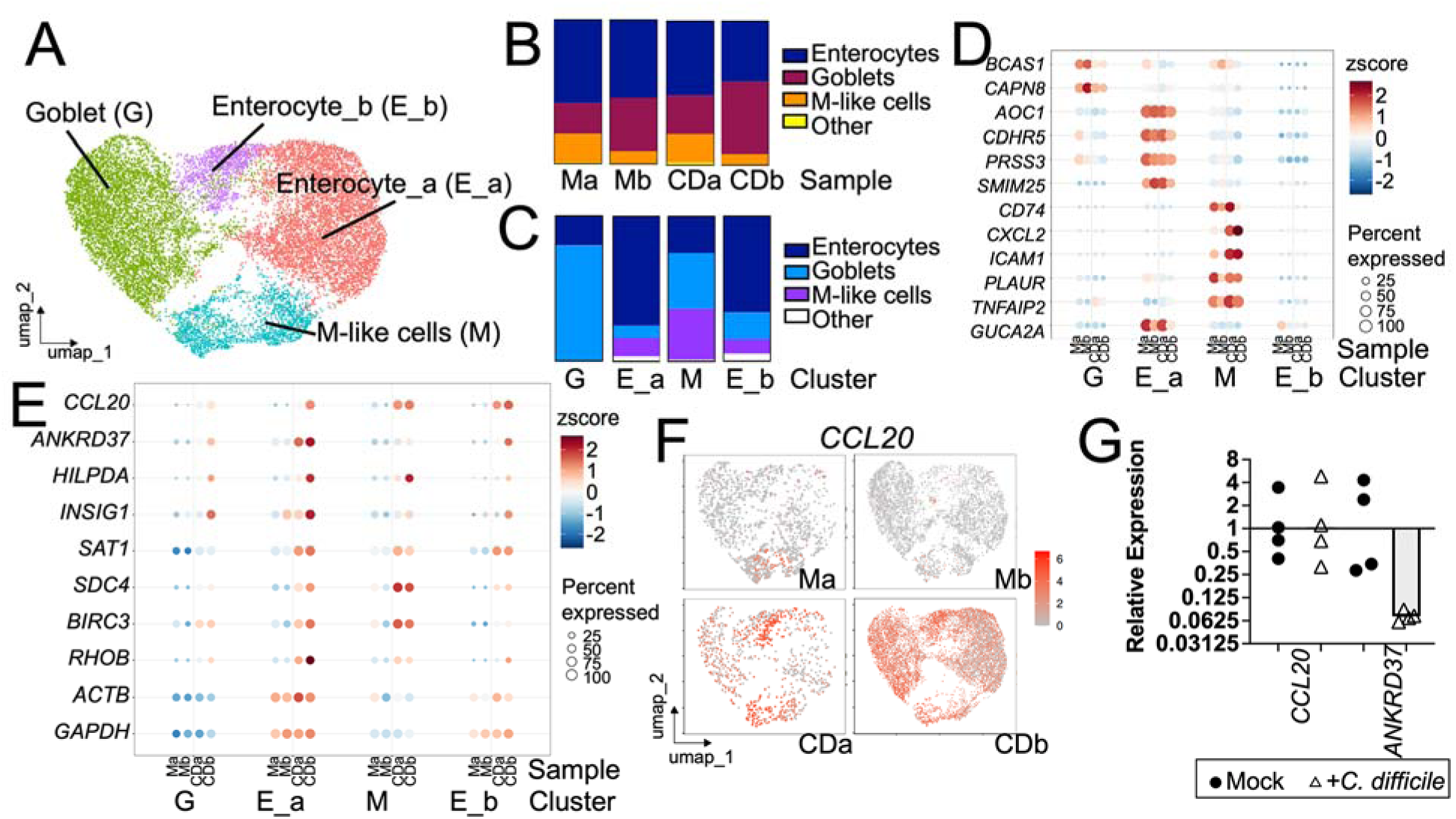
Markers expressed by colonoids during co-culture with C. difficile. Differentiated colonoid monolayers were produced as in Figure 1. Monolayers co-cultured with *C. difficile* or PBS for 15 hrs were harvested and subjected to single cell RNA-Seq (scRNASeq). Results were analyzed using Seurat as described in Materials and Methods. Panel A: Cells are shown in a UMAP organized based on transcriptional similarity. Clusters are indicated with colors and labeled based on the dominant cell type in the cluster as shown in panel C. Panel B: Cell type composition of each sample is shown. Ma, mock, replicate a; Mb, mock, replicate b; CDa, *C. difficile* co-culture, replicate a; CDb, *C. difficile* co-culture, replicate b. Panel C: Cell type composition of each cluster is shown. Other includes stem cells, TA cells and tuft cells. Panel D, Expression of cluster markers is shown in each cluster as a Dot Plot where the size of the dot indicates percent of cluster expressing the gene and the color indicates the z score. *BCAS1, CAPN8*, markers of goblet cells. *AOC1, CDHR5, PRSS3, SMIM24*, *GUCA2A*, markers of colonocytes. *CD74, CXCL2, ICAM1, PLAUR, TNFAIP2*, markers of Paneth-like cells/M cells. Panel E, Expression of genes is shown in each cluster as a Dot Plot where the size of the dot indicates percent of cluster expressing the gene and the color indicates the z score. *CCL20, ANKRD37, HILPDA, INSIG1, SAT1, SDC4, BIRC3* and *RHOB* are candidates for markers of colonoid response to *C. difficile* co-culture. *ACTB* and *GAPDH,* widely used housekeeping genes, are shown for comparison. Panel F, Feature plot indicating expression of *CCL20* in each cell in each sample. Grey indicates low expression, shades of red indicate higher expression. Panel G: Additional differentiated colonoid monolayers were produced as in Figure 1 and co-cultured with *C. difficile* (open triangles) or PBS (black circles) for 16 hrs. Medium was removed and monolayers were extracted with Trizol and the PureLink RNA purification kit as described in Materials and Methods. *CCL20* or *ANKRD37* gene expression normalized with *GAPDH* gene expression was measured in cDNA by real time RT qPCR. Results are shown relative to the geometric mean of mock samples. Each symbol shows results obtained with one monolayer. Bar, geometric mean.

To identify cell types, a reference-based method was employed using the SingleR package in R [36] with a reference based on an atlas of cells from healthy human colons [37]. The majority of the cells were annotated as enterocytes or goblet cells of varying maturity (Fig. 4BC). These results are consistent with immunofluorescence results (Fig. 1) showing that cells positive for NHE3, a marker of colonocytes, or MUC2, a marker of goblet cells, were readily detected. Some cells were annotated as more rare cell types including M cells, transit-amplifying cells, stem cells and tuft cells. Each sample contained the major cell types detected (Fig. 4B). Clusters were given names based on their most abundant predicted cell type (Fig. 4A,C). Cluster M was composed of a mixture of cells that were labeled enterocytes, goblets and M-like cells, and was named based on its most distinctive cell type.

Analysis of genetic markers for each cluster provided further support for the cell type labels. Genes expressed more highly in each cluster (cluster markers) relative to cells in all other clusters were identified. Some of the cluster markers were also previously described as markers of colonic cell types [34]; expression of a subset of these genes is shown in Fig. 4D (*BCAS1, CAPN8*, markers of goblet cells; *AOC1, CDHR5, PRSS3, SMIM24, GUCA2A*, markers of colonocytes; *CD74, CXCL2, ICAM1, PLAUR, TNFAIP2*, markers of cells labeled “Paneth-like cells” by Gao et al [34] or M cells by Smillie et al [37]). For most markers, cluster-specific upregulation in all samples was observed. Cells of cluster E_b did not show strong expression of cluster specific markers despite being mostly labeled as Enterocytes.

To identify markers of colonoid responses to *C. difficile* in co-culture, differential gene expression analysis comparing Mock and Cd samples in each cluster was conducted using FindMarkers() with log-fold-change threshold set to 0.25 and minimum fraction of cells expressing a gene at 0.1. Results were ranked by adjusted p-value and the top candidate gene for every cluster was *CCL20* (Supplemental Table S1). Markers that were expressed more highly in Cd samples and were among the top 50 candidate genes (based on adjusted p-value) in at least 3 of the 4 clusters were identified. Expression of these candidate genes is shown in Fig. 4E. *RHOB* was included because it was upregulated in 2/4 clusters and had been previously identified as differentially expressed in response to *C. difficile* [15, 17, 40, 41]. Widely used housekeeping genes *ACTB* (encoding Actin) and *GAPDH* were included for comparison (Fig. 4D). *CCL20* showed strong upregulation in response to *C. difficile* in both samples and upregulation was detected in cells of all clusters (Fig. 4D,E). *ANKRD37, HILPDA, INSIG1, SAT1, SDC4, BIRC3* and *RHOB* showed upregulation in multiple clusters. Additionally, there were some genes that exhibited upregulation in one or few clusters only. For example, *CXCL8,* encoding IL-8, was upregulated most consistently in cluster M (Table S1). Thus, several candidates for *C. difficile*-responsive genes were identified and *CCL20* was the top candidate. *CCL20* expression has been previously detected in response to *C. difficile* bacteria [24], supporting the results of this analysis.

Expression of *CCL20* and *ANKRD37* was further tested in colonoid monolayers by real time RT qPCR to determine whether upregulation could be detected using bulk RNA. Monolayers were produced and co-cultured with or without *C. difficile* for 16 hrs as above. RNA was extracted and real time RT qPCR was conducted as described in Materials and Methods. As shown in Fig. 4G, increased expression of *CCL20* or *ANKRD37* was not detected. The discrepancy between the results of the scRNASeq experiment and the analysis of bulk RNA may reflect heterogeneity in the responses of individual cells and the fact that the transcriptomes used for scRNASeq were heavily filtered (>250 and < 5000 detected expressed genes and < 40% mitochondrial expressed genes) so that only the most healthy cells were included in the analysis. In contrast, bulk RNA extraction included all cells in the monolayer, regardless of their health.

### CCL20 production during *C. difficile* co-culture detected by immunofluorescence

To confirm the results of the scRNASeq and determine whether CCL20 was useful as a marker of colonoid responses to *C. difficile* in co-culture, we employed immunofluorescence. Mock and *C. difficile* co-cultured colonoids were incubated for 16 hrs. CCL20 is a secreted protein and therefore, monolayers were treated with Brefeldin A (25 ug/ml final concentration) for 4 hrs to block protein secretion and enhance the ability to detect cells producing CCL20. Monolayers were washed, fixed with 4% PFA and probed with anti-CCL20 antibody, anti-*C. difficile* antibody, and Hoechst. Figure 5 A-D shows that bright CCL20 staining was observed in *C. difficile* co-cultures in contrast to mock inoculated cultures (Fig. 5, E-H). The overlays shown in Fig. 5 D,H include only the red channel (CCL20) and the green channel (*C. difficile*). These results thus demonstrated increased CCL20 production in *C. difficile* co-cultured colonoids, confirming the scRNASeq findings.

**Figure 5.**
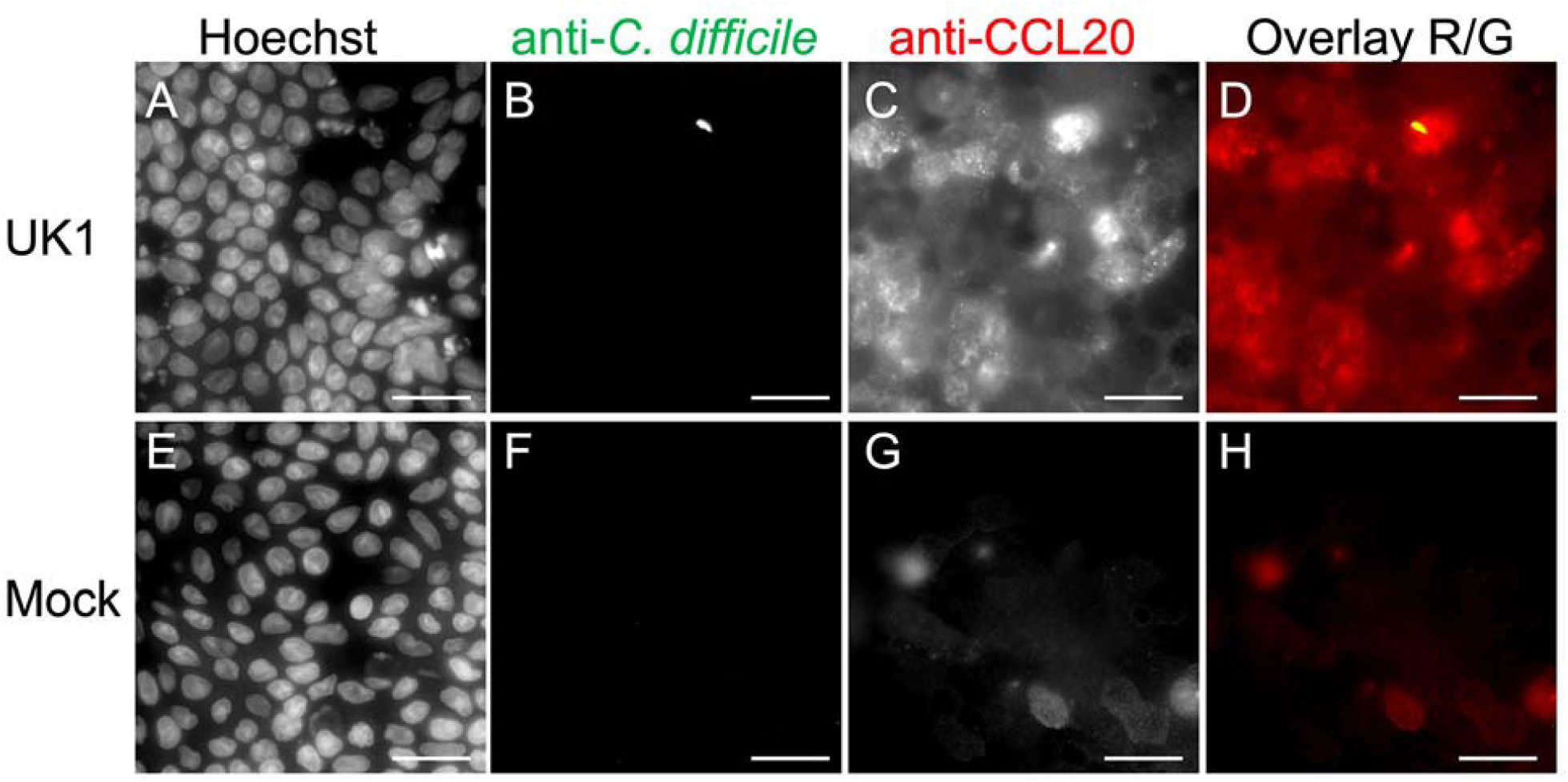
**Increased production of CCL20 in monolayers co-cultured with *C. difficile.*** Differentiated colonoid monolayers were produced as in Figure 1. Monolayers were inoculated with pre-germinated spores of *C. difficile* strain UK1 (panels A-D) or buffer (panels E-H) and co-cultured for 16 hrs. Brefeldin A (25 ug/ml final concentration) was added to the apical and basal media to block protein secretion and monolayers were incubated for an additional 4 hrs. Medium was removed and monolayers were washed, fixed in 4% PFA and stained for immunofluorescence. Panels A-H: staining with Hoechst (A, E); staining with anti-*C. difficile* (B, D (yellow), F, H); staining with anti-CCL20 (C, D (red), G, H (red)). Scale bar, 20 um. CCL20 production was observed in numerous cells throughout the co-cultured monolayer.

In some cases, CCL20-positive cells were located near a detected *C. difficile* bacterium (Fig. 5D). However, many responding cells lacked bacteria in their immediate vicinity at the time of imaging. Bacteria loosely associated with the monolayer may have been present during co-culture but were washed away during processing. Additionally, colonoids may be responding to secreted bacterial products such as toxins and thus nearby bacteria would not be required for CCL20 upregulation. The role of toxins in *C. difficile*-dependent production of CCL20 was therefore tested further.

### Non-toxigenic *C. difficile* have reduced ability to promote CCL20 production

As discussed above, the secreted glucosylating toxins, TcdA and TcdB, are major virulence factors of *C. difficile*. Therefore, to identify the contribution of these toxins to bacterial/colonoid interactions, we examined the behavior of nontoxigenic *C. difficile* in the colonoid co-culture system. TcdA and TcdB are encoded in the pathogenicity locus along with the sigma factor TcdR, which is required for toxin expression [9]. The nontoxigenic R20291::*tcdR* mutant [27] was compared to its toxigenic parent strain R20291 (an epidemic strain closely related to UK1 [26]).

Transwells containing differentiated colonoid monolayers were produced as described above and inoculated with either R20291, R20291::*tcdR* or buffer. These transwells were inoculated with vegetative cells because spores of strain R20291::*tcdR* show an altered response to germinants [27]. After 17 hrs of co-culture, the apical medium was collected, used to wash the monolayer, diluted and plated to enumerate *C. difficile* CFU as described in Materials and Methods. Vegetative cells of both strains grew in the colonoid co-culture system, reaching levels of 2-4 x 10^7^ CFU/transwell (Fig 6 I). Glucosylating toxin production was thus not required for bacterial growth in this system although the *tcdR* mutant strain reached a level approximately 2-fold lower than the WT strain. The *tcdR* mutant did not exhibit a defect in growth or saturation density under anaerobic conditions [27].

**Figure 6.**
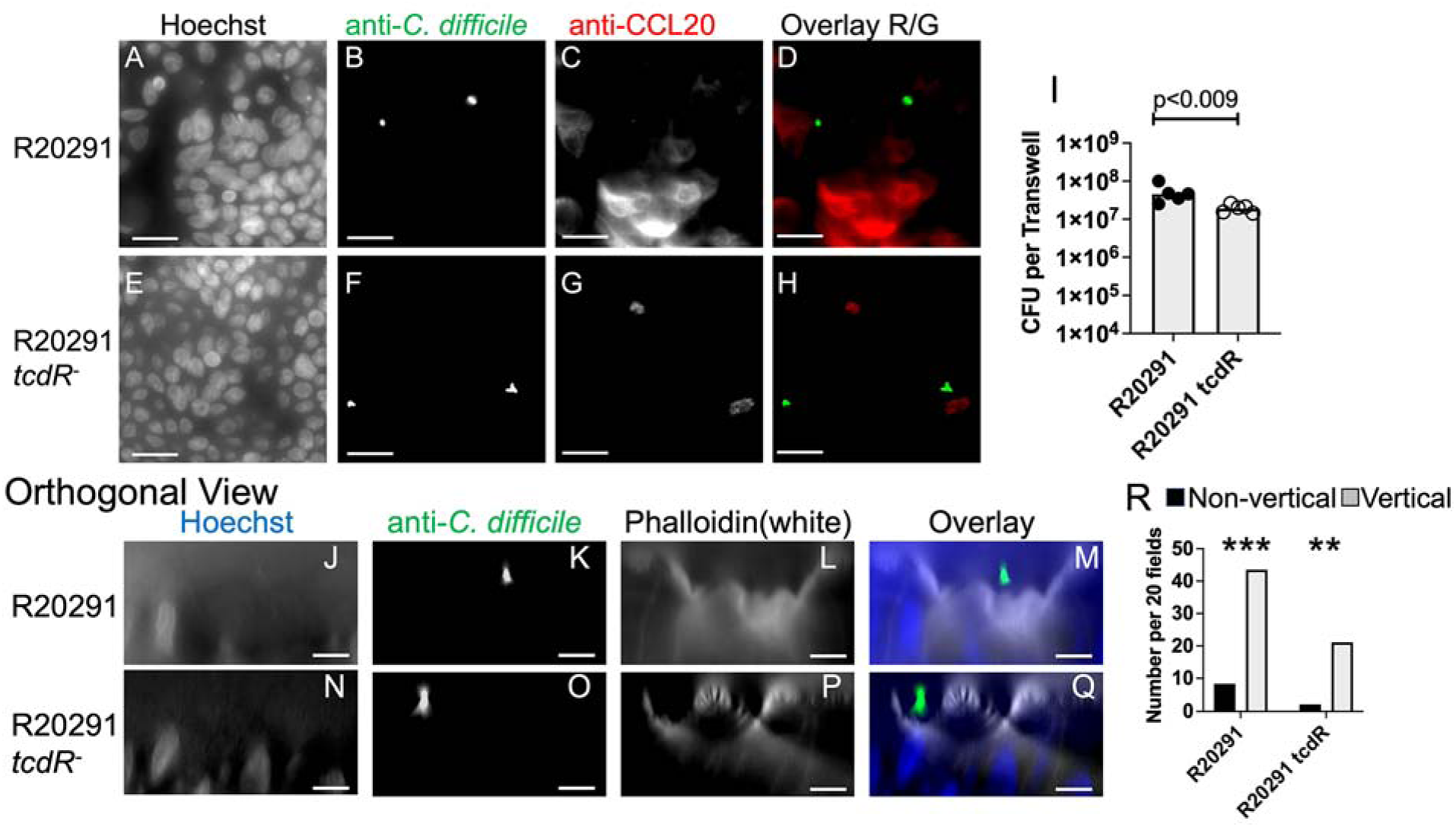
Reduced production of CCL20 by monolayers co-cultured with nontoxigenic *C. difficile*. Differentiated colonoid monolayers were produced as in Figure 1. Panels A-H: Monolayers were inoculated with vegetative cells of strain R20291 (A-D) or R20291 *tcdR* (E-H) and co-cultured for 16 hrs. Brefeldin A (25 ug/ml final) was added to the apical and basal media and monolayers were incubated for 4 hrs. Medium was removed and monolayers were washed, fixed in 4% PFA and stained for immunofluorescence. Panels A-H: staining with Hoechst (A, E); staining with anti-*C. difficile* (B, D (green), F, H); staining with anti-CCL20 (C, D (red), G, H (red)). Scale bar, 20 um. Panel I: Apical culture medium was collected in an anaerobic chamber and total CFU/transwell was determined. Each symbol shows results from a different transwell. Bar, geometric mean. p values, *t* test performed with log transformed data. Composite results from 2 experiments. Panels J-O: Monolayers were inoculated with vegetative cells of strain R20291 (J-M) or R20291::*tcdR* (N-O) and co-cultured for 17 hrs. Brefeldin A was not used. Monolayers were washed, fixed with 4% PFA, permeabilized and stained. Orthogonal views are shown with apical surface above and basal surface below. Stained with Hoechst (J, M (blue), N, Q (blue)); stained with anti-*C. difficile* (K, M (green), O, Q (green)); stained with phalloidin to stain filamentous actin (L, M (white), P, Q (white)). Scale bar = 10 um. Panel R: Bacterial orientation was scored as in Figure 3 and numbers were normalized to 20 microscope fields. Grey bars show vertical bacteria; black bars show non-vertical bacteria. Results tested against a 50:50 distribution by Fisher’s exact test. **, p < 0.004; ***, p<0.0004.

To test for CCL20 upregulation in colonoids, monolayers were co-cultured with R20291 or R20291::*tcdR* for 16 hrs. Brefeldin A was added to the apical and basal medium and the monolayers were incubated for 4 more hours. Monolayers were washed, fixed, permeabilized and incubated with antibody or stains for fluorescence microscopy. Colonoids co-cultured with WT R20291 showed expression of CCL20 (Fig 6 A-D) while colonoids co-cultured with R20291::*tcdR* did not show this response (Fig 6 E-H). The overlays shown in Fig. 6 D,H include only the red channel (CCL20) and the green channel (*C. difficile*). Thus, nontoxigenic bacteria did not promote production of CCL20 to the same extent as WT bacteria. These observations suggest that secreted toxins play a role in activating widespread responses to *C. difficile* in the colonoid monolayer system. Consistent with these results, previous studies demonstrated CCL20 upregulation by purified TcdB or *C. difficile* culture supernatants [24, 42].

Despite the failure to promote CCL20 production, non-toxigenic *C. difficile* strains associated with the monolayer in a vertical orientation. Monolayers were co-cultured with bacteria, with or without Brefeldin A, and then washed, fixed, permeabilized, stained and imaged. Phalloidin staining of filamentous actin was used to identify the apical surface of the cells. As shown in Fig 6 J-R, most of the monolayer-associated bacteria were in the vertical orientation for both strains. Bacteria also showed a vertical orientation after Brefeldin A treatment. Thus, toxin production and early colonoid responses to *C. difficile* challenge were not required for bacteria to associate with the monolayer in the vertical orientation.

## DISCUSSION

This study demonstrates that early interactions between *C. difficile* and the intestinal epithelium can be modeled in a simple tissue culture system using human colonoids. We found that *C. difficile* was able to grow in co-culture with colonoid-derived epithelial monolayers in standard tissue culture conditions (5% CO2, with ambient oxygen). In this simple system, bacterial association with the monolayer was detected after prolonged co-culture. Further, colonoid responses to *C. difficile*, such as upregulation of CCL20, were observed at a time point when only small numbers of tightly associated bacteria were detected with the monolayer. Early upregulation of CCL20 may represent a response to production of glucosylating toxins by *C. difficile* whereas bacterial association with the colonoid monolayer occurred independently of toxin production.

The observation that bacterial cells associated with the colonoid monolayer is consistent with previous studies demonstrating that *C. difficile* expresses proteins with adhesion functions [43]. For example, cell wall proteins Cwp2 [44] and Cwp66 [45] promote adherence to epithelial cells in vitro and CD2831 [46] and CbpA [47] bind components of the mammalian extracellular matrix such as collagen. These adhesion-associated proteins may contribute to binding of *C. difficile* bacteria to the colonoid monolayer during co-culture. The vertical orientation of bacteria detected in this study may indicate that this orientation promotes tighter association of bacteria with cells.

CCL20 was detected as a protein produced by colonoids in response to *C. difficile*. Previous studies have described similar upregulation of CCL20 in response to *C. difficile*. Live *C. difficile* bacterial culture or bacterial culture supernatant [24] or purified *C. difficile* toxin TcdB and flagella [42] upregulated CCL20; the effects of toxin and flagella were synergistic. In addition to CCL20, several other genes including *ANKRD37, HILPDA* and *INSIG1* were identified as *C. difficile* toxin-responsive, differentially expressed genes in previous transcriptomic studies [15–17, 40, 41, 48]. The results reported here are consistent with these earlier observations and further demonstrate that toxins produced by live bacteria are sufficient to stimulate this response.

Additionally, a variety of other conditions have been shown to upregulate *CCL20* in host cells. Several pathogens including *Salmonella typhimurium, Mycobacterium avium paratuberculosis, Salmonella enteritidis, Listeria monocytogenes* [24, 49], bacterial components such as flagellin [49] and Hepatitis C virus [50] increase CCL20 production.

CCL20 upregulation has also been detected in Inflammatory Bowel Disease (IBD) [51, 52]. IBD is a chronic condition characterized by abnormal inflammatory responses to normal microbiota. The responses evoked by excessive CCL20 production play an important role in this disease.

Only a small number of tightly associated bacteria were observed after 15 hrs. of co-culture. However, widespread responses to *C. difficile* were identified based on gene expression (Fig. 4D,E) and detection of CCL20 (Fig. 5). Notably, cells lacking any visibly adjacent bacteria also upregulated CCL20. This response was not triggered by non-toxigenic *C. difficile* (Fig. 6E-H). Together, these results argue that secreted toxins act both locally, near toxin producing bacteria and also at a distance, enabling widespread response throughout the monolayer. Given that bacterial populations exhibit heterogeneous expression of *tcdA* during infection [53], such widespread effects may help sustain the survival of bacterial cells that are not actively expressing toxin.

Previous studies have investigated *C. difficile*/epithelial cell interactions in tissue culture systems. One approach employs specialized apparatus to create an anerobic environment in the apical compartment and aerobic conditions in the basolateral compartment thereby supporting simultaneous growth of bacteria and epithelial cells [10–13, 54].

Simpler systems have co-cultured intestinal epithelial cells with *C. difficile* using short term anaerobic conditions [20] that are nonoptimal for the epithelial cells or high MOI under aerobic conditions [55] that are nonoptimal for the bacteria. To bypass the need for live bacteria entirely, other studies have examined the effects of individual *C. difficile* components such as TcdA [17], flagella [18], or membrane vesicles [19].

In the present study, we exploited the ability of colonoids growing in a transwell monolayer to support the growth of *C. difficile* in the apical medium above the monolayer. This co-culture system utilizes standard transwell inserts which are widely available and do not require specialized equipment to use. Advances in organoid culture have enabled major breakthroughs in understanding the pathogenicity of viruses and parasites that were previously uncultivable in tissue culture systems [56, 57]. Likewise, for *C. difficile*, the colonoid-based co-culture system offers considerable advantages for dissecting *C. difficile* virulence mechanisms, probing human epithelial responses, testing therapeutic interventions and performing high resolution imaging of the bacterial-epithelial interactions. Together, these benefits will help accelerate research into *C. difficile*-human epithelium interactions.

## CONCLUSIONS

This communication describes a co-culture system in which the strict anaerobe *C. difficile* proliferated with differentiated human colonoid monolayers derived from colonic biopsies and used to model the intestinal epithelium. Analysis of epithelial responses demonstrated upregulation of the chemokine CCL20 as a prominent early response to infection. These findings establish a human epithelial infection model and identify CCL20 activation as an early host response to *C. difficile* in this model, providing a framework for investigating mechanisms of CDI pathogenesis.

## ACKNOWLEDGMENTS

The authors thank Drs. Mary Estes, Linc Sonenshein, Boris Belitsky, Laurent Bouillaut, Shonna McBride, Aimee Shen, Ralph Isberg, Joan Mecsas, Alyssa Fasciano, Wai-Leung Ng, David Kaplan, Ying Chen and Terrence Roh for generously providing helpful suggestions and expertise. We also thank Emma Hayes for technical support. The authors acknowledge the Tufts University High Performance Compute Cluster, which was utilized for some of the research reported in this paper. All writing was human generated.

## DATA AVAILABILITY

scRNASeq data are deposited in the NCBI GEO database (accession number GSE327570).

## SUPPLEMENTARY MATERIALS

Figure S1: Mitochondrial gene expression in samples. Table S1: *C. difficile* responsive genes.

## AUTHOR CONTRIBUTIONS

Paola Zucchi and Adrianne D. Gladden contributed equally to this work. Conceptualization, P.Z. and C.A.K.; methodology, P.Z.; formal analysis, A.D.G, A.T., R.B. and C.A.K..; investigation, P.Z., A.W.D., J.D., and C.A.K..; resources, R.G., M.A., and J.R..; writing—original draft preparation, P.Z., A.D.G., and C.A.K.; writing—review and editing, R.G., A.W.D., J.D., M.A., J.R., A.T., and R.B.; supervision, C.A.K..; project administration, C.A.K.; funding acquisition, C.A.K.

