## Supplemental Figure S1 for "*Clostridioides difficile* stimulates *CCL20* expression in human colonoid monolayers in a transwell-based co-culture system that supports its anaerobic growth"

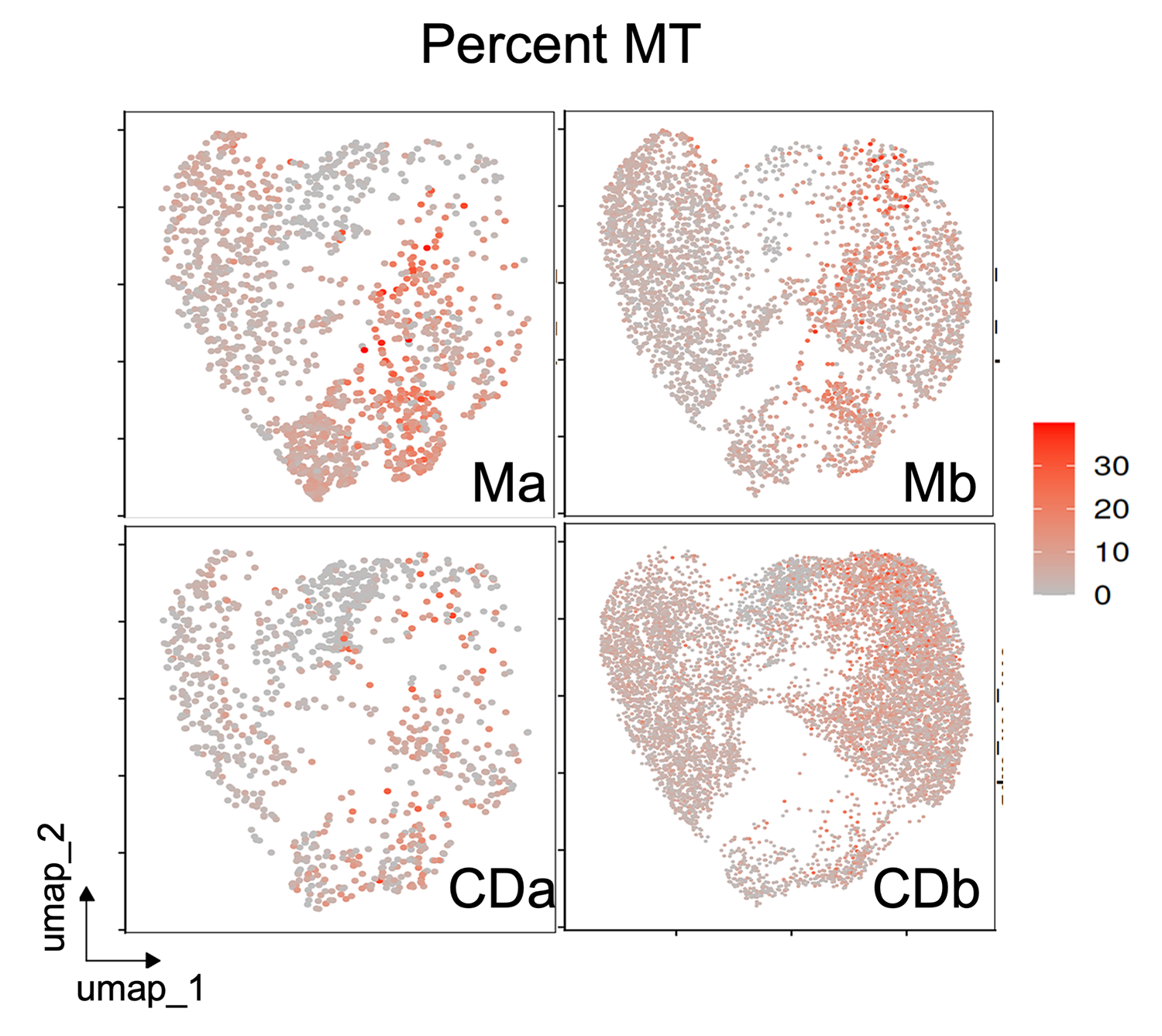

Supplemental Figure S1. Mitochondrial gene expression in samples.

Feature plot indicating percent mitochondrial gene expression (percent MT) in each cell in each sample. Grey indicates low percentage, shades of red indicate higher percentage.
